# Competence of DNA Tetrahedron and Exosomes as Nanocarriers for Epirubicin drug delivery in Breast Cancer Cells

**DOI:** 10.1101/2025.02.11.637654

**Authors:** Afnan Saleem, Sahar Saleem Bhat, Ubaid Gani, Nausheen Shafi, Payal Vaswani, Lateef A Dar, Junaid Nazir, Sheikh F Ahmad, Dhiraj Bhatia, Syed Mudasir Ahmad

**Affiliations:** Division of Animal Biotechnology, Sher-e-Kashmir University of Agricultural Sciences & Technology, Kashmir, India; University of Kashmir, India; Department of Biological Sciences and Engineering, Indian Institute of Technology Gandhinagar, Palaj, Gujarat, India; Department of Pharmacology and Toxicology, College of pharmacy, King Saud University, Riyadh-11451, Saudi Arabia

**Keywords:** DNA tetrahedron, drug delivery, exosomes, breast cancer, epirubicin, metastasis

## Abstract

Epirubicin, a chemotherapeutic agent, is used in treatment of metastatic breast cancer treatment. However, poor intracellular delivery and systemic toxicity are its major limitations. Our study evaluates the use of DNA tetrahedron (TD) structures and exosomes for targeted delivery of epirubicin to enhance its therapeutic efficacy. Increased intracellular concentration of epirubicin was observed in Flow cytometer when delivered via the TD: Epi system compared to free epirubicin. Cell viability assays using Annexin V-FITC and propidium iodide staining revealed higher apoptosis rates in the TD: Epi system. Exosomes, primarily 100 ± 20 nm in size (99% of particles), exhibit a - 28.5 mV zeta potential, ensuring colloidal stability. SEM and TEM confirm their spherical shape, uniform size, and double membrane structure. The TD: Epi system showed the highest efficacy in RT-qPCR analysis, followed by Epirubicin + Exosomes systems and epirubicin alone, further corroborated with western blot results. While exosomes also provided improved delivery and therapeutic outcomes compared to free epirubicin, they were less effective than the TD: Epi system. These results suggest that the TD carrier significantly enhances the delivery and apoptotic efficacy of epirubicin when compared to exosome-based delivery or epirubicin alone offering a promising strategy for improving treatment outcomes.

**Graphical Abstract:** 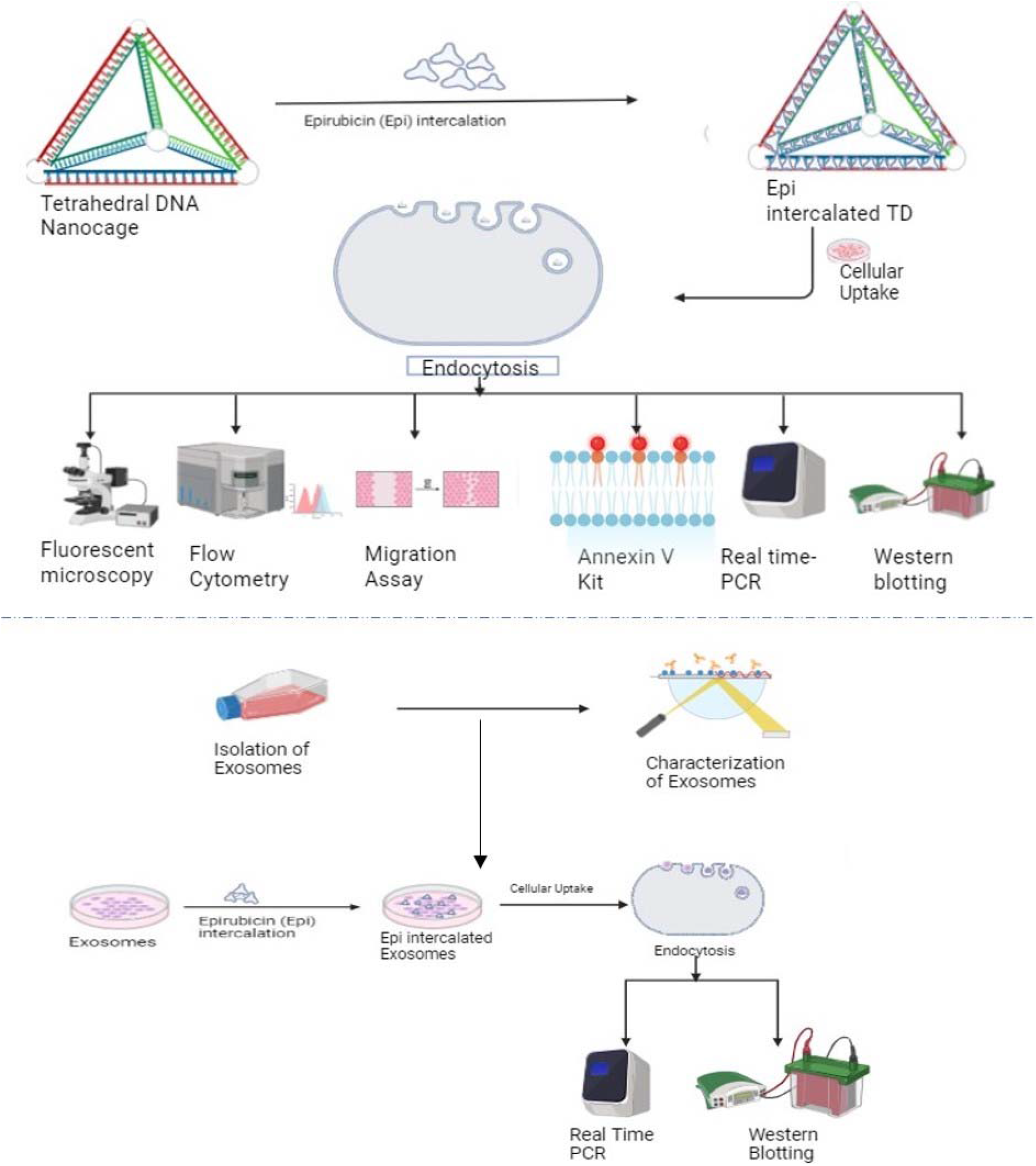

## Introduction

Breast cancer is the second leading cause of death for women globally and the most prevalent cancer diagnosed in women (Watkins 2019). Based on molecular subtypes, it can be classified as triple negative breast cancer (TNBC), human epidermal growth factor receptor 2 (Her 2) positive, luminal A, and luminal B. These differ greatly in terms of prognosis, treatment approaches, and metastasis (Jin and Mu 2015). Estrogen receptor (ER), progesterone receptor (PR), and Her2 receptor are not expressed in TNBC. 15 – 20% of breast cancers accounts for TNBC which is usually identified at an advanced diagnosis stage. It results in a high recurrence rate and low survival rate (Garrido-Castro, Lin, and Polyak 2019; Dent et al. 2007). Numerous FDA-approved drugs have been developed and used for patients, yet they continue to have various limitations. Major conventional treatment for metastatic cancer include surgery, hormonal therapy, radiation, chemotherapy and immunotherapy. Introducing targeted treatment can potentially reduce mortality and improve quality of life in patients with breast cancer. Epirubicin is an anthracycline and an epimer of doxorubicin with important roles in early and metastatic breast cancer treatment. Epirubicin efficacy is similar to doxorubicin, however, they have a different toxicity profile in regard to cardiotoxicity (Khasraw, Bell, and Dang 2012). Anthracycline chemotherapy is a standard practice for breast cancer and several other malignancies. Epirubicin is the most commonly used anthracycline in combination with other chemotherapy regimens for breast cancer, achieving higher survival than other alternatives (Joshi and Press 2018). Epirubicin acts as a topoisomerase II inhibitor, causing DNA damage and acute oxidative stress in cells which in turn triggers mitochondria mediated apoptosis (Loibl and Gianni 2017).

However, despite being an efficient chemotherapeutic agent, the drug’s myocardiopathy and tumor cells resistance are its main side effects. Targeted delivery of this drug through nanocages helps to overcome such side effects (Yazdian-Robati et al. 2022). A three-dimensional pyramidal nanostructure is the tetrahedral DNA nanostructure (TDN) which is formed by the complementary pairing of four single-stranded DNA. It is being recommended as a viable drug carrier due to its great stability, biocompatibility, exceptional cellular uptake rate, abundance of functional modification sites, and adaptability for different drugs (Goodman, Berry, and Turberfield 2004; Goodman et al. 2005).

Exosomes are extracellular vesicles (40-160 nm) which are secreted by live cells. They can be also found in different biological fluids (e.g., urine, saliva, and serum). Small size, low toxicity, good biocompatibility, and low immunogenicity are reported advantages of using exosomes as drug delivery vehicles. Drug-loading strategies include physical loading techniques (e.g., electroporation, ultrasound and extrusion), incubation, and cell engineering techniques etc. (Zhang et al. 2023). Exosomes as drug delivery vehicles holds significant promise in targeted therapy for metastatic breast cancer.

In this study, we engineered a novel drug delivery system using DNA nanocages in the form of tetrahedrons to encapsulate epirubicin drug. These nanocages are meticulously designed to provide a stable and controlled environment for drug delivery, ensuring targeted release and enhanced therapeutic efficacy. Our approach leverages the structural precision of DNA nanotechnology to achieve efficient loading and release kinetics, promising significant advancements in cancer treatment strategies. We also evaluated the intrinsic ability of exosomes as natural carriers and facilitating efficient drug delivery to cancer cells. We compared the DNA tetrahedron and exosomes for targeted deliver of epirubicin in comparison to free epirubicin in metastatic breast cancer treatment.

## 2. Materials & methods

### 2.1 Tetrahedron (TD) synthesis

The one-pot technique was used to synthesize TD. To put it briefly, 2 mM MgCl2 was mixed with the four oligonucleotides (T1, T2, T3, and T4) in equal molar ratios (1:1:1:1). In a thermocycler, the reaction mixture was first heated to 95 °C, then annealed and cooled gradually to 4 °C. This cooling process was done in 5 °C decrements, with a 15-minute interval at each step. A final TD concentration of 2.5 µM was attained.

### 2.2 Synthesis of TD: Epi

Epirubicin (100 μM) and tetrahedron (1 μM) were incubated in a shaker incubator (350 rpm) at 37°C for three hours. Unloaded epirubicin was removed by centrifugation at 10,000rpm for 10 minutes. Following the centrifugation, supernatant was collected. TD: Loaded cages were desalted to remove any unbound drug.

### 2.3 TD and TD: Epi system characterization

#### 2.3.1 Electrophoretic mobility shift assay (EMSA)

To confirm the formation of a higher order structure, EMSA was performed using 10% Native-PAGE. Sample contained 10 μl of TD: Epi, 6 μl of 1X TAE, and 3 μl of loading dye. As a control, we also loaded genomic DNA incubated with nanocage & nanocage only in subsequent wells. The gel was run at 100V for 60 min. The gel was stained with EtBr stain and a Gel Documentation system was used to visualize it (BioRad ChemiDoc MP Imaging system).

### 2.4 Cell culture studies

Human breast cancer cell lines MDA-MB-231 and MCF-7 were procured from the National Centre for Cell Science (NCCS) Pune. Cells were cultured in DMEM media (Sigma-Aldrich, St Louis, MO, USA) which was supplemented with 10% fetal bovine serum (FBS) (Sigma-Aldrich) and 4 mM L-glutamine (Invitrogen, Waltham, MA, USA). A humidified incubator at 37°C with 5% CO_2_ was used to maintain the cells.

### 2.5 Cellular uptake

Breast cancer cell lines (MDA-MB-231 & MCF-7) were seeded in culture plates (6-well) and allowed to grow till 80% confluency. DPBS washings were given to cells and serum starved for 20 mins. Cells were then treated with epirubicin only (100 μM) and TD: Epi system (1:100) for 1 hour in serum free-media at 37°C. Untreated wells were kept as controls. After 1-hour incubation, media was decanted and cells were washed with DPBS twice to remove any unbound system. Cells were stained with DAPI and the uptake was then analyzed with fluorescent microscope (Axiovert).

### 2.6 Scratch assay

MDA-MB-231 & MCF-7 cell lines were seeded in culture plates (6-well) and allowed to grow till confluency (100%). Using a standardized 10 μl tip, a scratch was created at the center of the wells in epirubicin only and TD: Epi system (1:100) wells. Untreated wells were kept as controls. The cells were washed with 1X DPBS to remove the detached cells. Cells were then incubated with fresh serum-free media with epirubicin only and TD: Epi system (1:100). The Axiovert microscope was used to observe cells at 10X at various time intervals (0, 6, and 24 hours). Fiji Image J software was used to process the images. The wound closure was assessed to measure the distance between the scratch.

### 2.7 Flow cytometry

Cells (MDA-MB-231 & MCF-7) were seeded in a culture plate (6-well) and allowed to grow till confluency (80%). The cells were rinsed with 1X DPBS and serum starved for 20 minutes. The cells were then treated with epirubicin alone and TD: Epi system (1:100) for 1 hour in serum free-media at 37°C. Untreated wells were kept as controls. After 1-hour incubation, media was decanted and cells were washed with 1X DPBS twice to remove unbound system. Cells were then trypsinized, the pellet was collected and dissolved in filtered DPBS. Cells were subjected to BD LSRFortessa X-20 Flow Cytometer to analyze the cellular uptake.

### 2.8 Fluorescent microscopy for apoptotic cells

Cells (MDA-MB-231 & MCF-7) were seeded in a culture plate (6-well) and allowed to grow till confluency (80%). Cells were washed with DPBS and serum starved for 20 mins. Cells were then treated with epirubicin only and TD: Epi system (1:100) in serum free-media at 37°C for 1 hour. Untreated cells were kept as controls. After 1-hour incubation, media was emptied and cells were washed with DPBS twice to remove unbound system. The detection of apoptosis was done using the Dead Cell Apoptosis Kit with Annexin V FITC & Propidium Iodide for fluorescent microscopy (Invitrogen) (Cat No: V13242). After the incubation period, cells were harvested and washed in cold DPBS. Following this, cells were centrifuged and the supernatant was discarded. Cells were then resuspended in1X annexin-binding buffer. The cell density was determined (∼1 × 10^6^ cells/ml) and diluted in 1X annexin binding buffer. 5 - 25 μl of the Annexin V conjugate and 1–2 μl of the 100 μg/mL PI working solution was added to each 100 μl of cell suspension. A sufficient volume was prepared per assay for deposition on a clean slide. Slides were mounted and observed under the fluorescence microscope using appropriate filters. Dead cells will show strong nuclear staining from PI and membrane staining by annexin V. However, live cells will show weak annexin V staining, while the apoptotic cells show high degree of surface labelling.

### 2.9 Exosomes isolation & characterization

#### 2.9.1 Isolation of exosomes

Bone marrow mesenchymal stem cells (hMSCs) (Cat No: PT2501) were grown till passage 3 and 80-90% confluency. Cells were rinsed thrice with sterile DPBS and cultured in serum-free media at 37 °C & 5% CO_2_ for 48 hours. Conditioned media was collected in 50 ml falcon tubes. Exosomes were isolated by modified differential centrifugation and ultracentrifugation on Ultracentrifuge-T865 Rotor Head (Thermo Scientific). Tubes were centrifuged at 300g for 10 mins to pellet down the cells. The clear supernatant was transferred to a fresh 50 ml tube. The fresh tubes were again centrifuged for 20 min at 2000 × g, at 4 °C to remove the large vesicles and dead cells. The clear supernatant was again transferred to a fresh tube and centrifuged at 1000g at 4 °C for 30 min to pellet down cell debris. Using a 0.22 µm sterile filter, supernatant was filtered and transferred into polyallomer tubes or polycarbonate bottles appropriate for the ultracentrifugation. Supernatant was again centrifuged to pellet down exosomes and unwanted proteins at 120,000 × g, at 4 °C for 70 mins. The supernatant was decanted carefully and the pellet was resuspended in 1 ml PBS in new polyallomer tube. The tubes were again centrifuged for 70 min at 120,000g at 4 °C. The supernatant was removed and exosome pellet was resuspended in small volumes of PBS (100-150 μL). Exosomes were stored at -80 °C till further use (Théry et al. 2006).

#### 2.9.2 Characterization of exosomes

##### 2.9.2.1 Scanning electron microscopy (SEM)

Scanning electron microscopy was used to determine the size and surface morphology of the isolated exosomes. 5 µl of MSC-exosome suspension was drop casted on plasma cleaned glass coverslips. Exosomes were fixed with 4 % paraformaldehyde solution and kept at 4 °C for 4 hour and vacuum dried overnight. A coating of 2 – 5 nm gold coating was applied by sputter coating technique and then imaged at an accelerated voltage of 10 kV by scanning electron microscope (Zeiss EVO 18). Exosome size analysis was done by ImageJ software, and the density distribution of exosome diameters were determined.

##### 2.9.2.2 Transmission electron microscopy (TEM)

The Transmission Electron Microscopy (TEM) analysis of exosomes was conducted by first fixing the exosomes (5 µL) in 1% glutaraldehyde on a formvar/carbon grid. Following fixation, the samples were counterstained with 1% uranyl acetate. The imaging was performed at an accelerating voltage of 120 kV to visualize the morphology and size distribution of the exosomes.

##### 2.9.2.3 Dynamic light scattering (DLS)

DLS was used to analyze the average diameter and particle number of exosomes. Exosomes were diluted to 0.1 mg/mL using Milli-Q. 200 µl was transferred into a cuvette. Size, vesicle population and zeta potential were measured using Malvern Zetasizer ZS90. The measurements were done in triplicates.

### 2.10 Co-incubation of exosomes & epirubicin

Drugs are loaded into exosomes using either the passive loading or incubation technique. This approach promotes drug integration into exosomes by leveraging the concentration gradient that arises when the external drug concentration is high. This approach effectively loads hydrophobic medicines onto exosomes by interacting with their lipid surface (Koh et al. 2023). Exosomes were incubated with epirubicin drug for 4 hours at 37°C. After incubation, the exosomes + epirubicin system was directly given to cells to evaluate the drug delivery efficacy of exosomes

### 2.11 Real-time expression

#### 2.11.1 RNA isolation & cDNA synthesis

RNA extraction was performed using Trizol method (Chomczynski and Sacchi 1987) from control, epirubicin only, TD: Epi system & Exosomes + Epirubicin system after 48 hr incubation in MDA-MB-231 & MCF-7 cell lines. The concentration and purity of the extracted RNA was measured in Nanodrop spectrophotometer (Thermo, USA). Complementary DNA was synthesized from isolated RNA using Revert Aid First Strand cDNA synthesis kit (Thermo Scientific) following the manufacturer’s instructions.

#### 2.11.2 Quantification by Real-Time PCR

Polymerase chain reaction (PCR) was carried out using primers mentioned in table 1. The 25µl reaction was carried out in a Veriti thermocycler (Applied Biosystems) with the following components. PCR cycling conditions included initial denaturation at 95°C for 5 min, denaturation at 95°C for 30 s, annealing at 60°C for 30 s, Extension at 72°C for 30 s, and final extension at 72°C for 5 min.

**Table 1:**
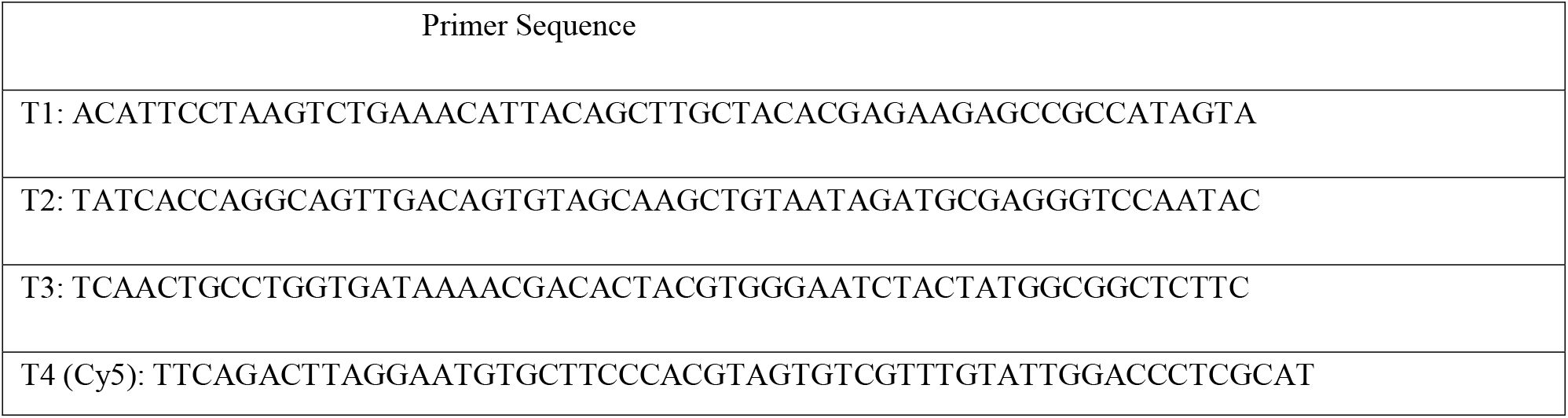
Tetrahedron Primer Sequence (5’ to 3’)

Real-time quantitative RT-qPCR was performed using SYBR Green PCR Master Mix (KAPA™ SYBR® qPCR Kit, Kapa Biosystems, Woburn, MA). Experiments were performed in duplicates. qPCR reactions were performed in QuantStudio 5 (ThermoFisher Scientific) and data normalized to GAPDH and β-Actin, which were used as internal controls. Primers used for expression were already reported (Table 2). GAPDH and β-actin were used as internal controls. Primers were purchased from Sigma-Aldrich.

**Table 2:**
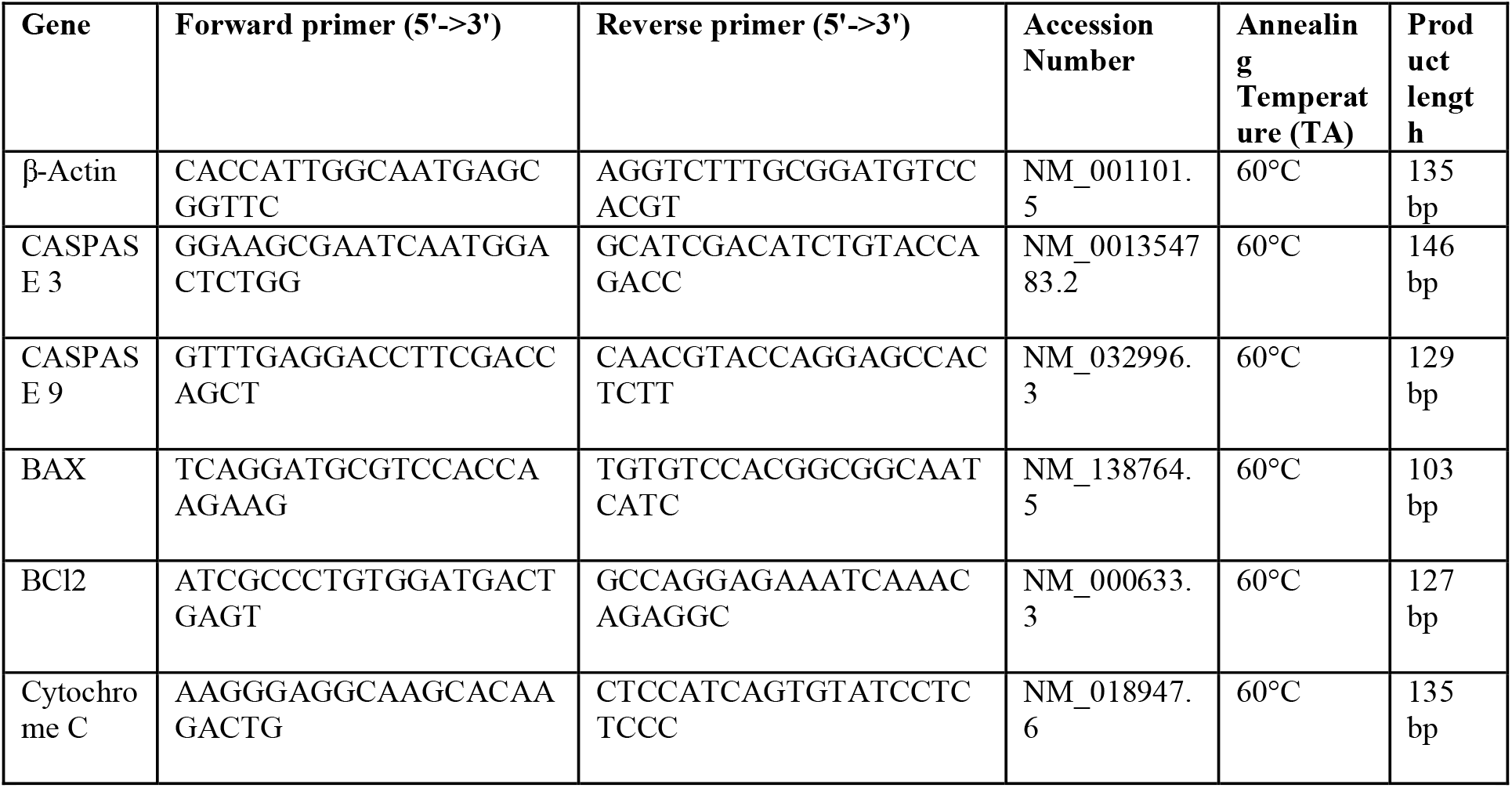
Primer details [FP (forward primer); RP (reverse primer); TA (Annealing temperature)].

#### 2.11.3 Statistical analysis

Data was represented as fold change in target gene expression, normalized to endogenous reference gene relative to calibrator using 2^−ΔΔCt^. GraphPad Prism 8.0 was used to perform the statistical analysis. Data is presented as mean ± standard error of the mean (SEM). Unpaired Student’s T-test and by one-way ANOVA was used to analyze the differences between means. Statistical significance was declared at P < 0.05.

### 3.0 Protein isolation and western blot analysis

Whole protein was extracted using RIPA lysis buffer supplemented with protease inhibitor cocktail (PIC) (#Cat No. P8340, Sigma-Aldrich), Phenyl methyl sulfonyl fluoride (PMSF) (#Cat No. RM1592, Himedia), and Sodium fluoride (NaF) (Himedia) from control, epirubicin only, TD; Epi system & Epirubicin + exosomes systems after 48-hour incubation in MDA-MB-231 & MCF-7 cell lines as previously described (Ali et al. 2022). Protein concentration was measured by Bradford’s method and absorbance was measured at 595 nm in a Spectrophotometer (Thermo Scientific). Readings obtained were plotted against a standard curve for BSA to retrieve the concentrations of protein. Western blotting was performed using Primary antibodies using caspase-3 (CST, Cat No. 14220, 1:5000) to probe protein bands. Secondary antibody used were (CST, S^3^ Rabbit, 1:3000). ChemiDoc MP (BioRad) was used to visualize the protein bands.

## 3.0 Results & Discussion

### 3.1 Synthesis & characterization of DNA nanocages (Tetrahedron)

Tetrahedron was synthesized using a previously described protocol (Goodman et al. 2005). Briefly, one pot synthesis method wherein all four primers were mixed in equimolar ratio was followed. It was subjected to thermal annealing from 95°C to 4°C. Final TD was characterized using electrophoretic mobility shift assay (EMSA). This thermal cycling promotes the precise folding and hybridization of the DNA strands, resulting in a stable tetrahedral configuration (Figure 1).

**Figure 1:**
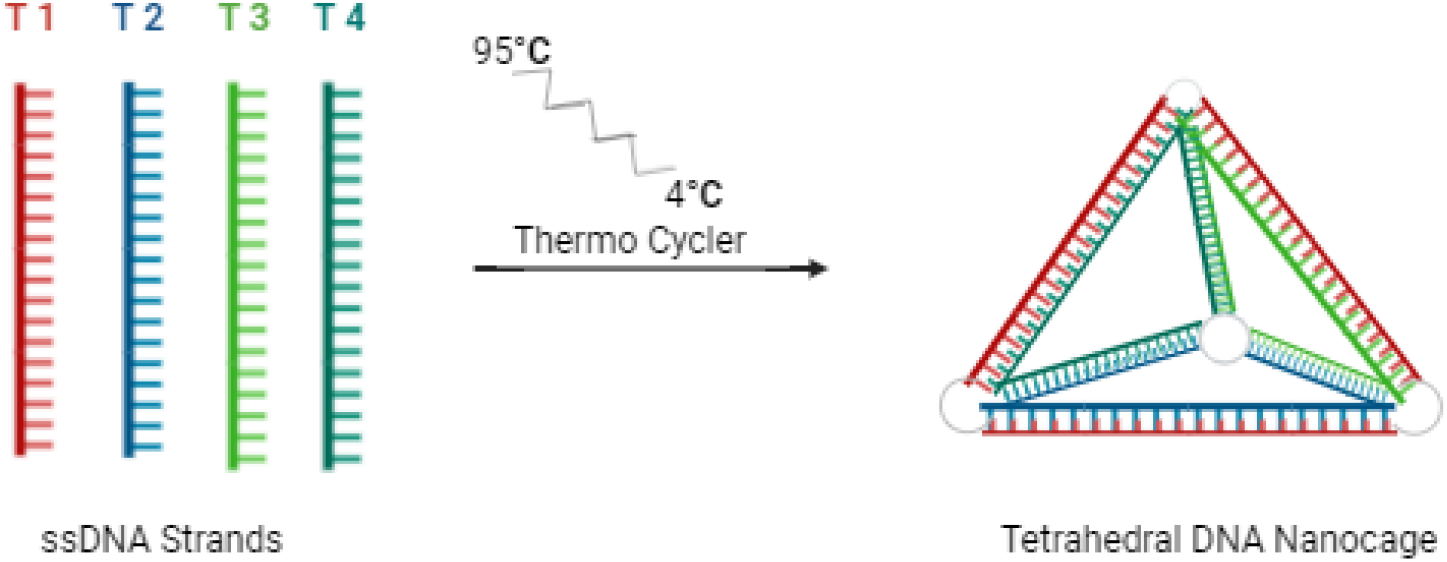
Synthesis of tetrahedron in a thermocycler: The four signal strands of DNA (ssDNA) were taken in equimolar ratio of 1:1:1:1. The reaction mixture was first heated to 95 °C, then annealed and cooled gradually to 4 °C. This cooling process was done in 5 °C decrements, with a 15-minute interval at each step.

Epirubicin drug was loaded onto synthesized TD nanocages. TD: Epi (1: 100 μM) was kept constant for further experiments. Loading was done at 37°C for three hours in a shaker incubator (350 rpm). Unloaded epirubicin was removed by centrifugation at 10,000rpm for 10 minutes. Following the centrifugation, supernatant was collected. Loaded cages were desalted to remove any unbound drug.

Synthesized TD: Epi was characterized using electrophoretic mobility shift assay (EMSA). Loaded cages were desalted to remove any excess unbound epirubicin. EMSA helps in detecting shift in the migration of DNA fragments on a gel due to changes in the mobility of DNA caused by drug binding on 10% native PAGE. When a DNA nanocage interacts with epirubicin drug, the complex formed migrates more slowly during gel electrophoresis compared to free DNA, resulting in a shifted band. The successful characterization of the synthesized TD: Epi system demonstrates the effective binding of epirubicin to the tetrahedral DNA nanocages. Desalting removed any excess unbound epirubicin ensuring that the observed shift in the gel electrophoresis were due to drug-DNA interactions. The distinct retardation of the bands on the 10% native PAGE correspond to the stoichiometric ratio of the oligos added (Figure 2). Ethidium bromide and epirubicin are capable of interacting with DNA and thus, TD; Epi system’s band intensity was lesser which confirmed the loading of epirubicin with TD. Since nanocages have been designed to deliver drug to cells, their behavior in EMSA alongside genomic

**Figure 2:**
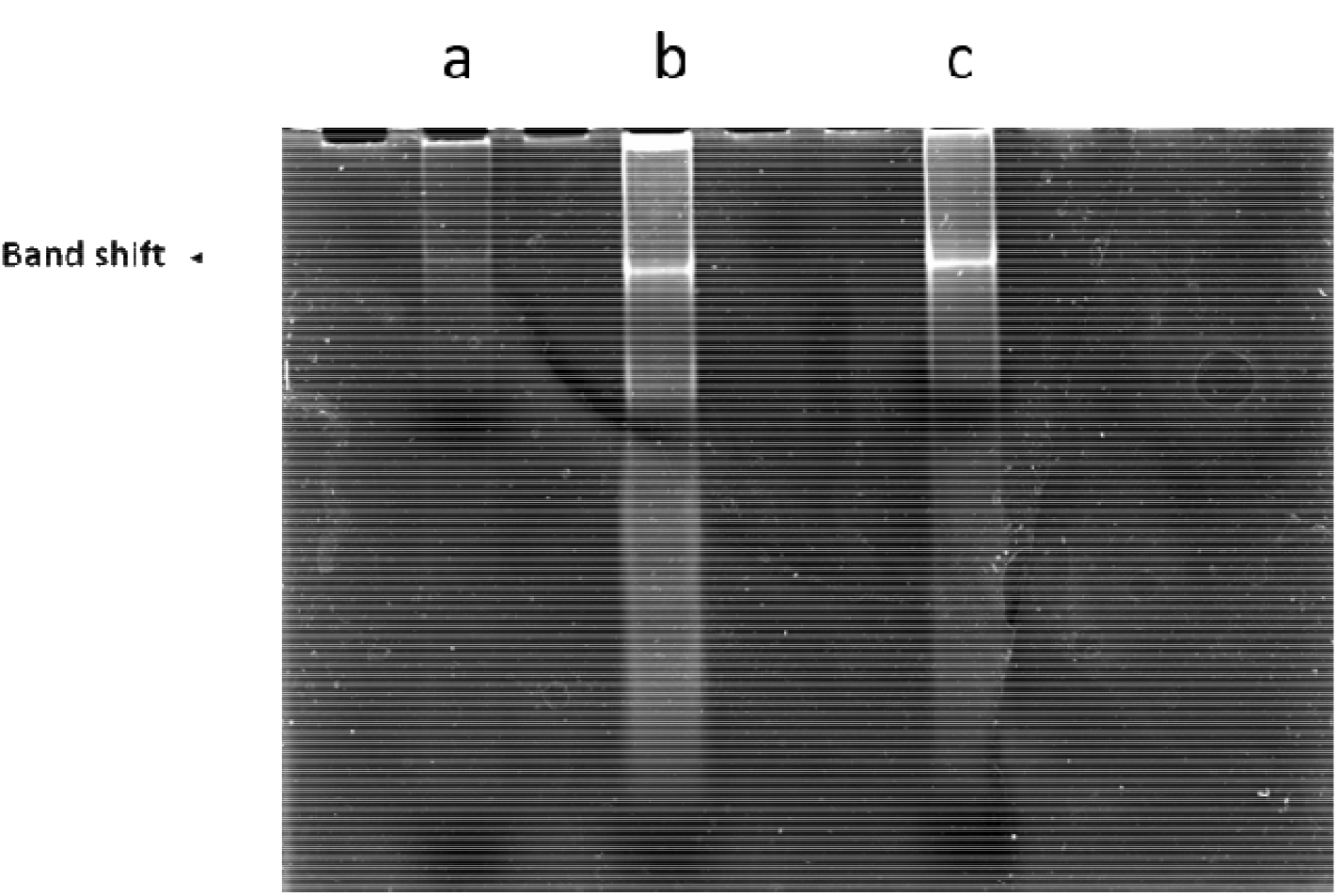
EMSA on 10% Native PAGE a) TD loaded with epirubicin (1: 100); b) Genomic DNA loaded with TD; c) Nanocage only.

DNA demonstrates their effectiveness and binding properties and thus, validates the functionality of the nanocages.

### 3.2 Cellular uptake using DAPI

MDA-MB-231 metastatic breast cancer cells were incubated with Epirubicin only and TD: Epi system for 1 hour to investigate if TDs enter the cells or not. The fluorescent intensity was quantified by using DAPI nuclear staining technique as DAPI binds to DNA (Figure 3). Fluorescent images confirmed the accumulation of epirubicin in the nucleus due to release of drug from TD: Epi system. Lowered fluorescence intensity observed in the TD: Epi system can be attributed to the more attractive delivery of epirubicin into the cells by the DNA nanocages. This provided clear evidence of epirubicin accumulation in the nucleus due to competitive binding of epirubicin to the DNA, displacing DAPI and thereby, reducing its fluorescent signal.

**Figure 3:**
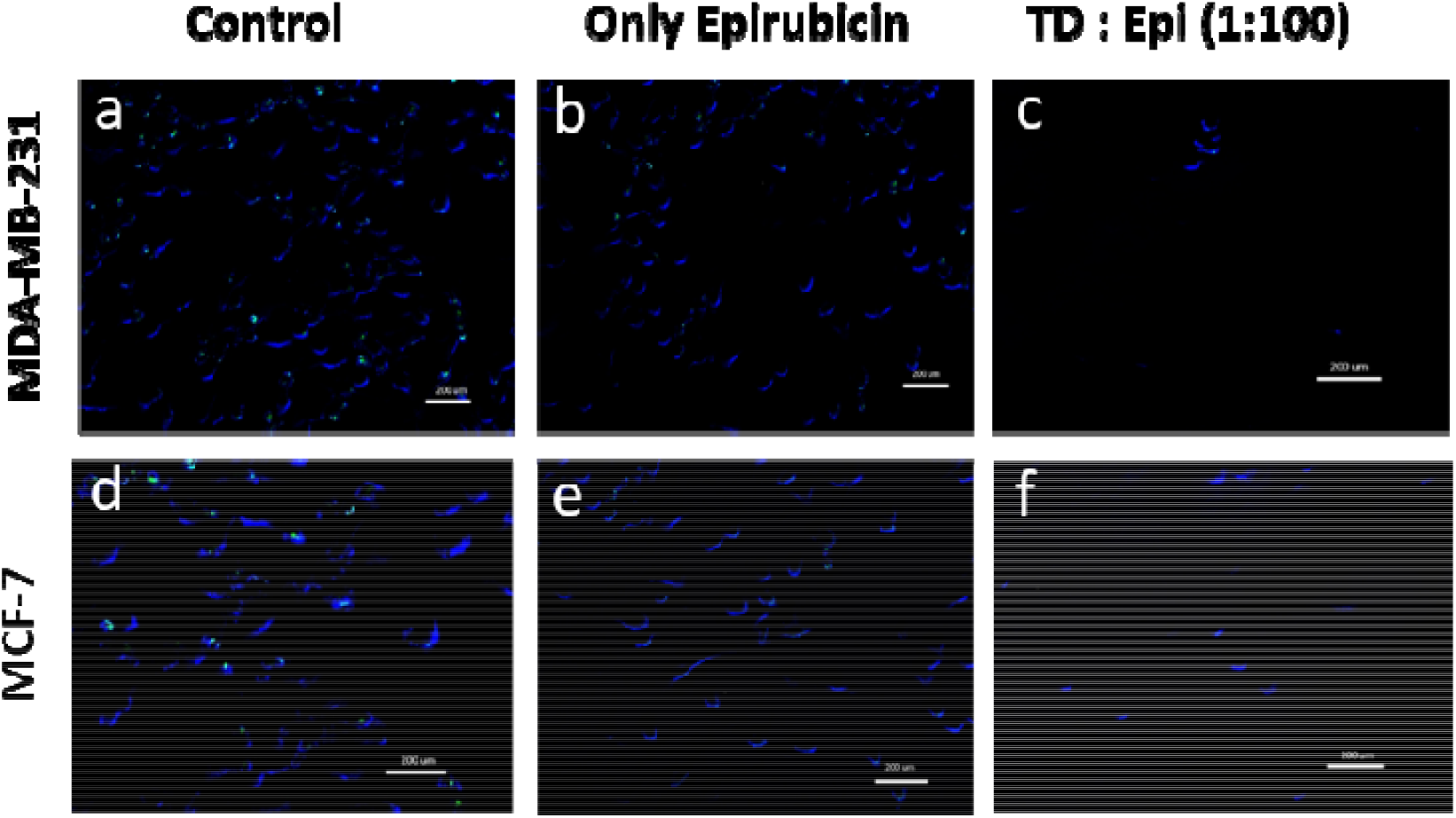
Accumulation of DAPI in MDA-MB-231 cells a) Control; b) Epirubicin only; c) TD: Epi (1: 100). Accumulation of DAPI in MCF-7 cells d) Control; e) Epirubicin only; f) TD: Epi (1: 100).

The study demonstrates TD: Epi system significantly enhances the intracellular delivery of epirubicin than epirubicin alone, resulting in higher nuclear accumulation and effective cytotoxicity against metastatic breast cancer cells.

### 3.3 Scratch assay

Scratch assay is a 2D cell migration assay wherein an artificial gap is formed and the movement is tracked using microscopy. In metastatic breast cancer cell lines, tetrahedron has been reported to promote cell migration (Gada et al. 2023). However, epirubicin shows a significant decrease in cell migration following exposure to 1 µM epirubicin (Wang et al. 2015).

In our study, we found that control & TD treated wells had good migration ability at all time intervals. However, epirubicin only & TD: Epi system had a reduced migration ability owing to the drug’s ability to inhibit migration & decrease the chance of metastasis. This effect is attributed to epirubicin drug’s ability to hinder cellular migration and hence reducing the metastasis potential of breast carcinoma cells (Figure 4).

**Figure 4:**
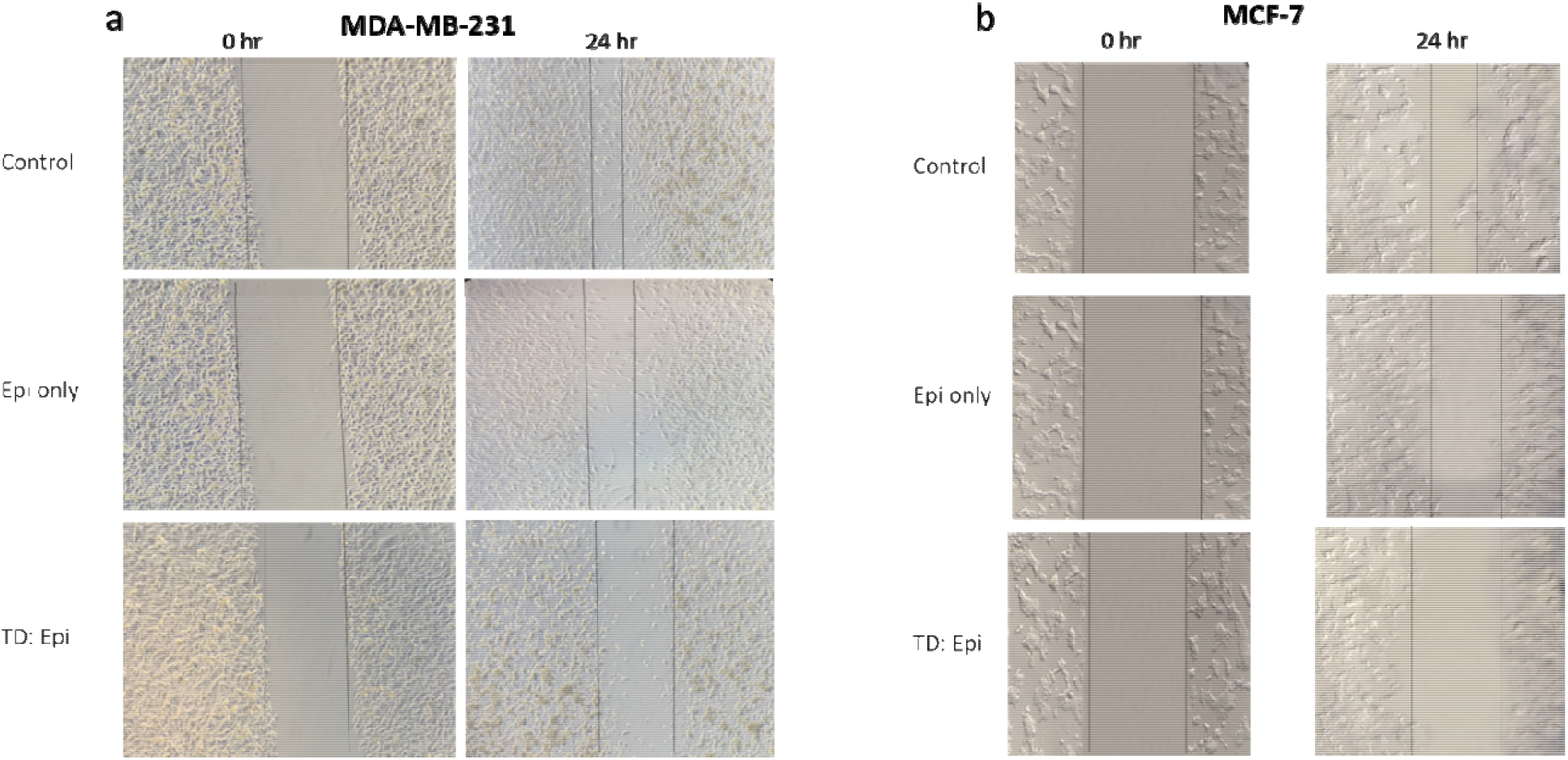
Scratch assay for a) MDA-MB-231; b) MCF-7.

Interestingly, when compared to free epirubicin, TD: Epi was found to be more effective in inhibiting the migration ability of cells. The enhanced effect suggests TD nanocarrier may facilitate more targeted and potent delivery of epirubicin, thereby exerting a stronger inhibitory influence on cancer cell migration.

### 3.4 Flow cytometry

Flow cytometry was performed to confirm the uptake of epirubicin & TD: Epi system in MDA-MB-231 cells for 1 hour. We acquired 10,000 events & gated them for epirubicin positive population. Results showed that TD: Epi had more amount of epirubicin accumulation inside the cells when compared to free epirubicin. The enhanced uptake suggests that the tetrahedron DNA carrier effectively enhances the intracellular delivery of epirubicin, potentially augmenting its therapeutic efficacy against metastatic breast cancer cells (Figure 5).

**Figure 5:**
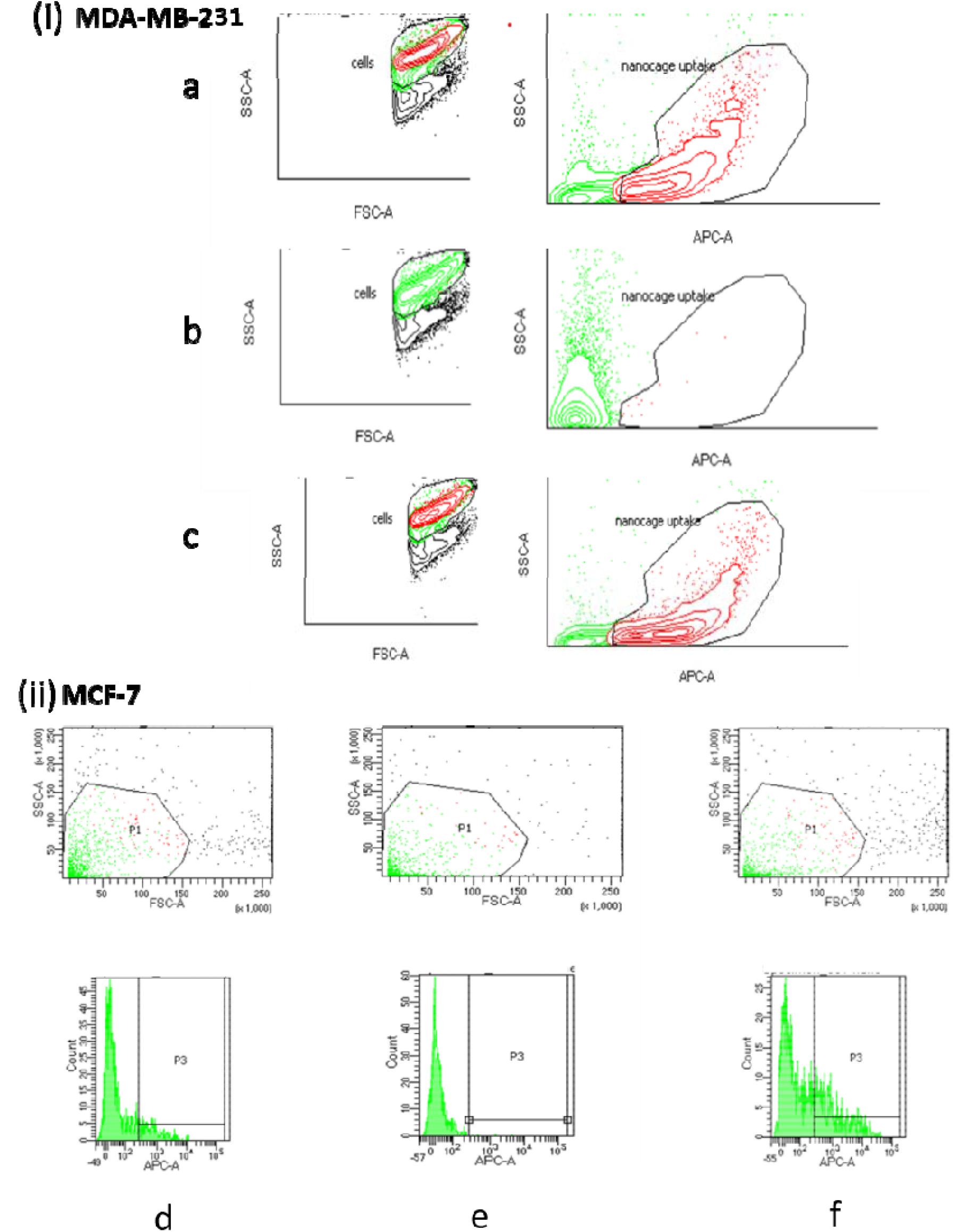
(i)Cellular uptake using flow cytometer in MDA-MB-231 a) TD: Epi system (1: 100); b) Epirubicin only; c) Nanocages only. (ii) Cellular uptake using flow cytometer in MCF-7 d) TD: Epi system (1: 100); e) Epirubicin only; f) Nanocages only

Our study found out when compared with epirubicin only, the fluorescence intensity of TD: Epi system was increased suggesting enhanced cellular uptake and efficient delivery of the drug by the TD nanocages. Epirubicin mechanism of action involves intercalating into DNA and inhibiting DNA helicase, thereby preventing replication and transcription (Townsend 2007).

### 3.5 Fluorescent microscopy for apoptotic cells

Propidium iodide and Annexin V-FITC were used to measure the cell viability. The staining method makes use of the characteristics of PI, which stains the DNA of cells that have lost membrane integrity and indicates late apoptosis or necrosis, and Annexin V-FITC, which binds to phosphatidylserine residues that have translocated to the outer leaflet of the plasma membrane early in apoptosis.

In this study, MDA-MB-231 cells were treated with Epirubicin only & TD: Epi system. Cells were subjected to Annexin V-FITC/PI staining to assess the cell viability. Cells that have bound Annexin V-FITC exhibited green staining in the plasma membrane while red staining (PI) throughout the nucleus was exhibited by those cells which have lost membrane integrity.

The fluorescent microscopy revealed increased presence of early apoptotic cells (green fluorescence) in the TD: Epi system indicates this delivery system initiates cell death more effectively and rapidly than free epirubicin (Figure 6).

**Figure 6:**
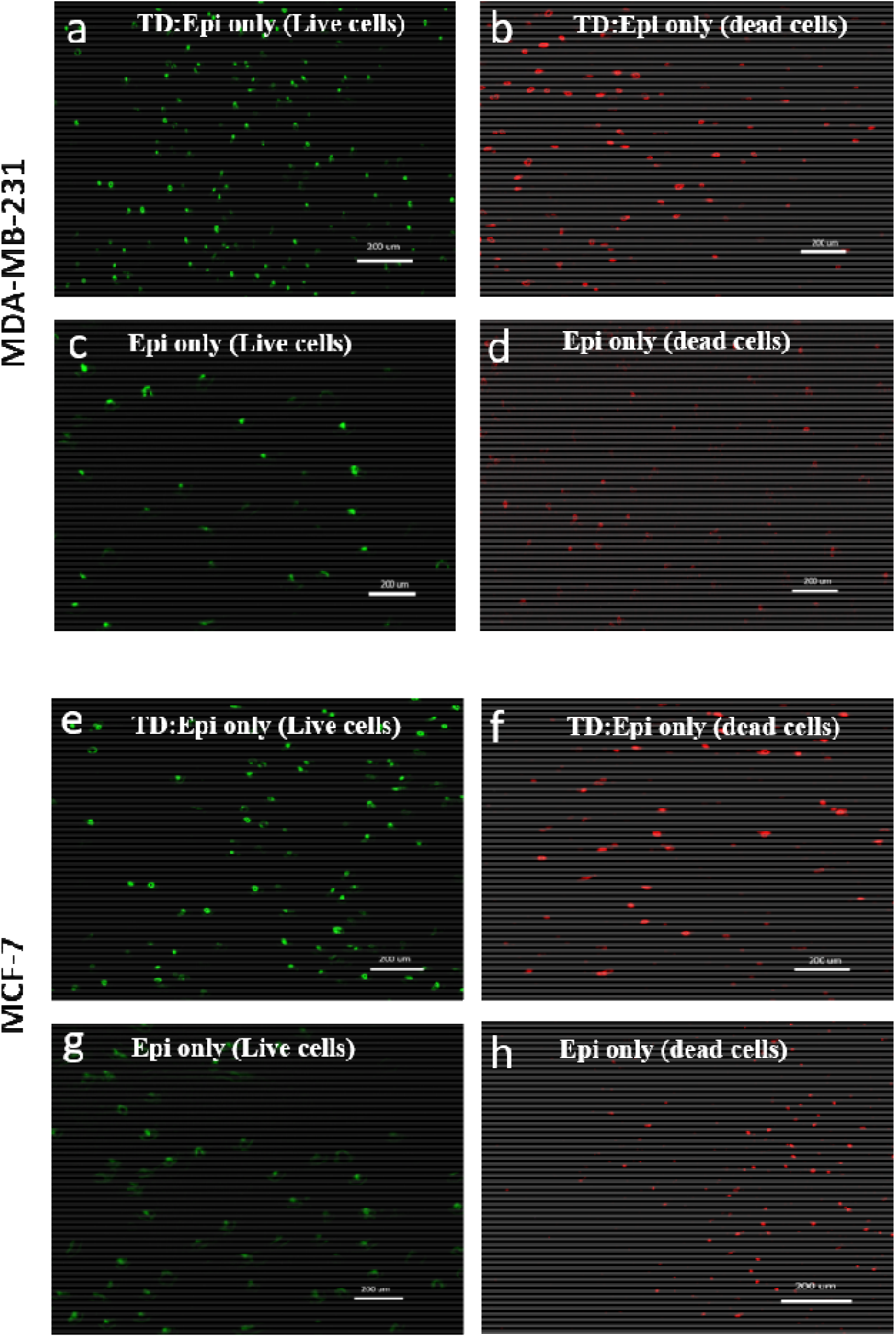
Fluorescent microscopy for live/dead cells in MDA-MB-231 cells and in MCF-7 cells a) TD: Epi (Live cells); b) TD: Epi (Dead cells); c) Epi only (Live cells); d) Epi only (Dead cells) ; e) TD: Epi (Live cells); f) TD: Epi (Dead cells); g) Epi only (Live cells); h) Epi only (Dead cells)

### 3.6 Exosomes isolation & Characterization

Exosomes are small extracellular vesicles, typically 30-150 nm in diameter, released by cells into the extracellular environment. Exosomes are being extensively studied for their potential in diagnostics, therapeutics, and regenerative medicine. The therapeutic efficacy of exosomes is largely attributed to the pro-migratory effects of their cargo, including growth factors, mRNAs and miRNAs, which significantly influence wound healing (Kalluri and LeBleu 2020; He et al. 2019). The DLS analysis shows the size distribution of the exosomes by the number of particles. DLS analysis indicated that the majority of particles (99%) were within the size range of 100 ± 20 nm. This confirms the homogeneity of the exosome population. The zeta potential measurements indicate the surface charge of the exosomes. The exosomes exhibit a zeta potential of -28.5 mV, suggesting good colloidal stability and a negative surface charge, which is crucial for maintaining exosome dispersion in biological fluids (Figure 7d). The SEM images reveal the surface morphology and particle size of the exosomes. The exosomes appear spherical with a uniform size distribution, confirming their nanoscale dimensions (Figure 7a). TEM imaging provides detailed visualization of the exosomes internal structure. The exosomes exhibit a characteristic double membrane vesicular structure and a distinctive cup-shaped morphology, as shown in the TEM images. (Figure 7b).

**Figure 7:**
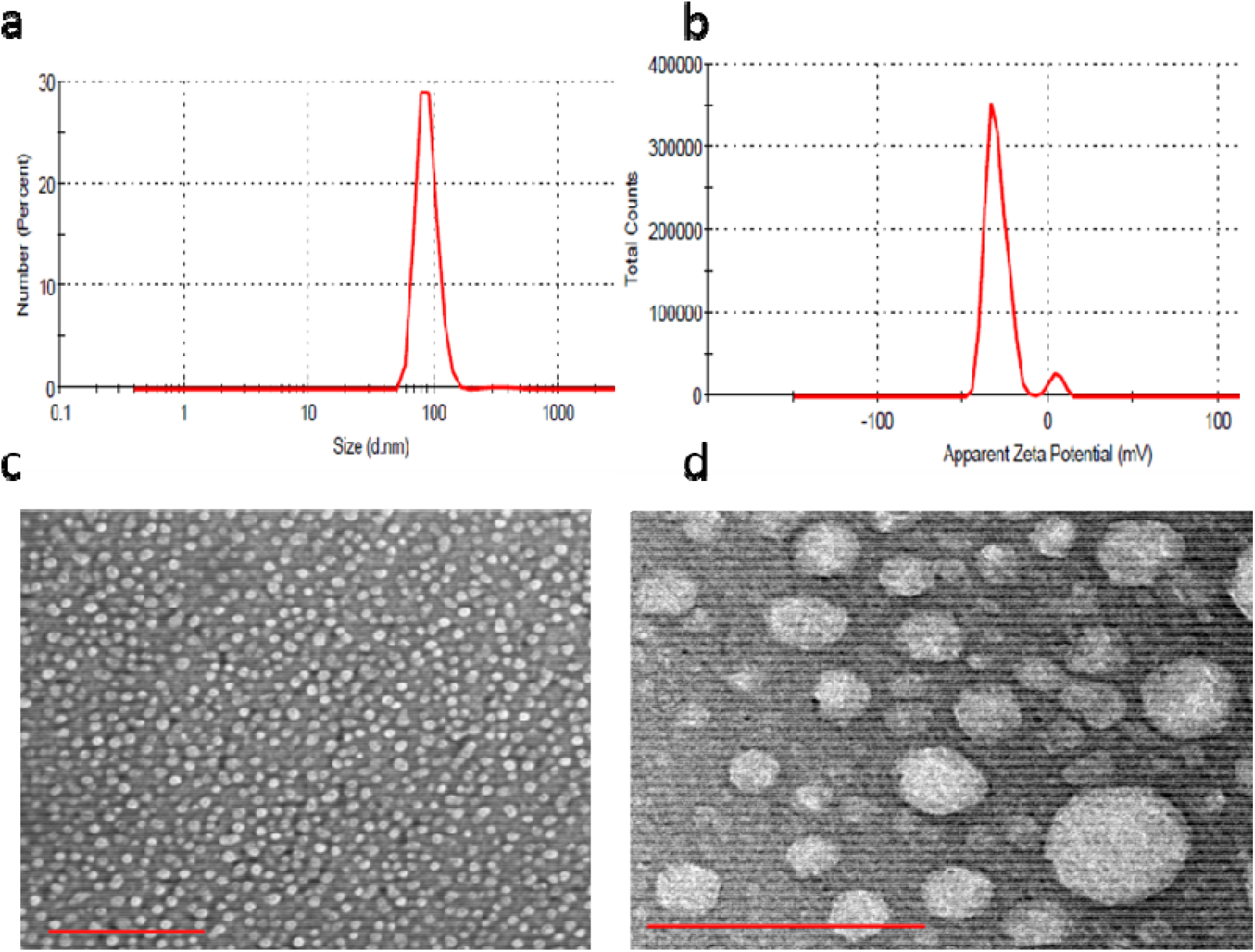
Characterization of bone marrow-derived stem cells (BMSCs) exosomes. (a) Dynamic light scattering (DLS) showing size distribution by number of particles, (b) zeta potential of exosomes, (c) Scanning electron microscopy (SEM) image showing surface morphology and particle size of exosomes; Scale bar: 200 nm (d) Transmission electron microscopy (TEM) image showing the double membrane vesicular structure of the exosomes with cup-shaped morphology. Scale bar: 100 nm.

### 3.7 RT-qPCR expression of apoptotic genes

mRNA expression of apoptotic genes (Caspase 3, Caspase 9, Cytochrome C, Bax and Bcl-2) was determined through RT-qPCR in control, epirubicin only, TD: Epi system and Epirubicin + Exosomes system in MDA-MB-231 & MCF-7 cell line (Figure 8). Our study showed significant upregulation of caspase 3, caspase 9, Cytochrome C and Bax and a significant downregulation of Bcl-2 gene in both the cell lines (Figure 9) indicating the activation of apoptotic pathway. The expression in TD: Epi system was the highest followed by epirubicin + exosomes system and epirubicin only indicating the higher efficacy of DNA tetrahedron as a carrier of epirubicin for breast cancer treatment when compared with epirubicin + exosomes system and epirubicin only.

**Figure 8:**
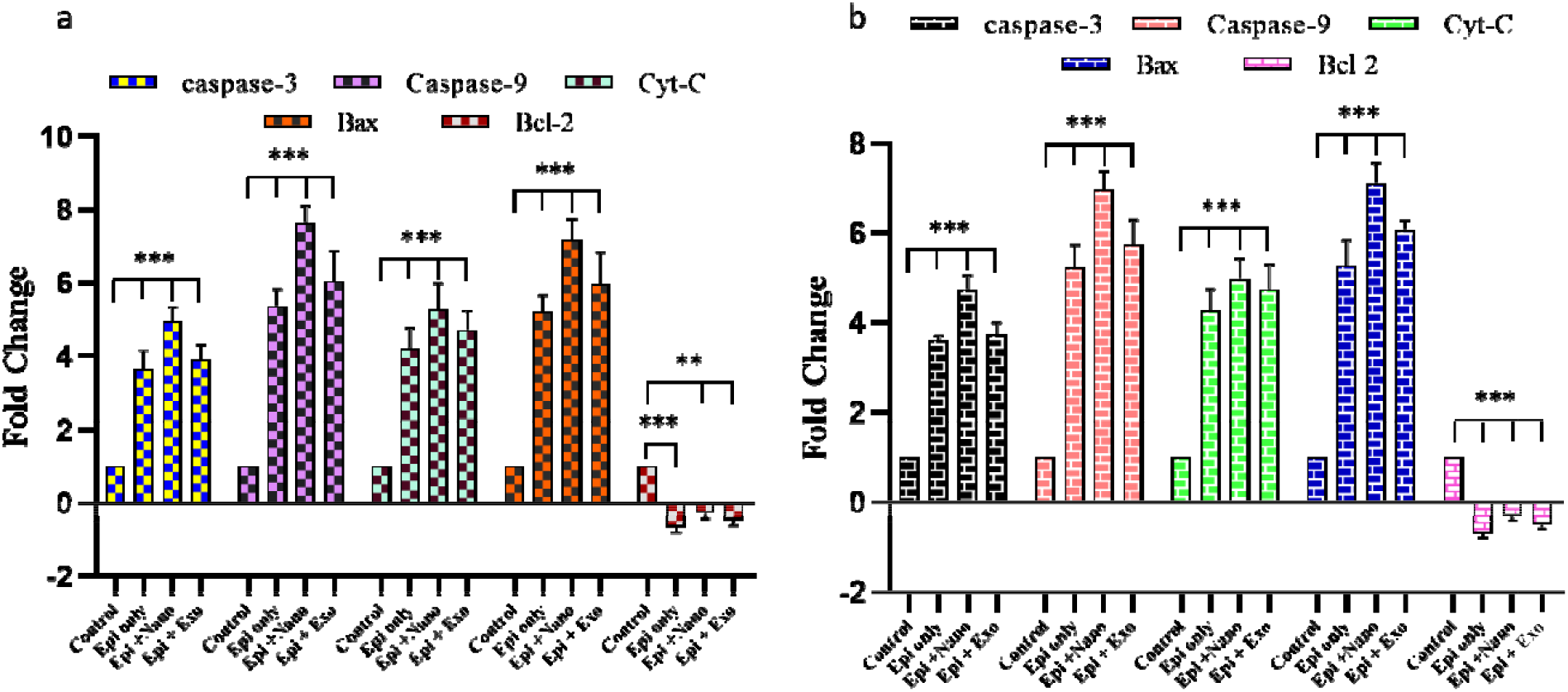
RT-qPCR expression of apoptotic genes a) MDA-MB-231; b) MCF-7.

**Figure 9:**
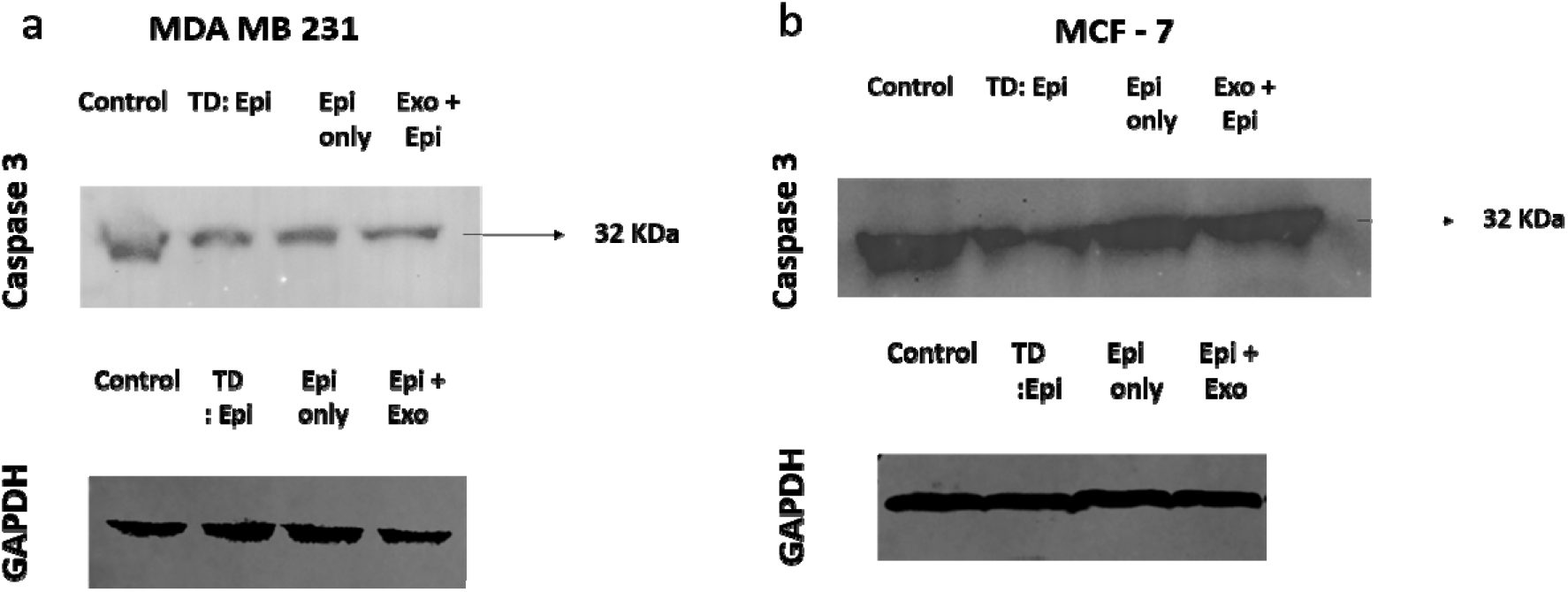
Western blot expression of a) Caspase 3 (MDA-MB-231); b) GAPDH (MDA-MB-231); c) Caspase 3 (MCF-7); d) GAPDH (MCF-7)

Specifically, TD: Epi system exhibited the highest levels of mRNA expression for Caspase 3, Caspase 9, Cytochrome C, and Bax, suggesting a robust activation of the intrinsic apoptotic pathway. Caspase 3 and caspase 9 are critical executors of apoptosis, while cytochrome C release from mitochondria is a key event triggering caspase activation. Bax, a pro-apoptotic member of the Bcl-2 family, promotes mitochondrial outer membrane permeabilization, further facilitating Cytochrome C release. Significant downregulation of Bcl-2 protein inhibits Bax and stabilized the mitochondrial membrane and thus, enhancing the pro-apoptotic environment induced by TD: Epi system.

Epirubicin + exosomes and epirubicin only and showed upregulation of pro-apoptotic genes, but to a lesser extent than the TD: Epi system. This led us to the conclusion that while both treatments are effective in inducing apoptosis, the TD: Epi system is a more potent system as it not only improves the intracellular delivery of epirubicin but also amplifies its apoptotic impact on cancer cells.

### 3.8 Protein expression of Caspase 3

Our study demonstrated a marked decrease in the protein expression levels of pro-caspase 3 in TD: Epi system followed by epirubicin + exosomes system and epirubicin alone in MDA-MB-231 & MCF-7 cell lines (Figure 9). Caspase 3 is a key executioner of apoptosis. This suggests that the apoptosis was induced in our experimental model contributing to the maximum overall cell death observed in TD: Epi system indicating its efficacy as a carrier of epirubicin for breast cancer treatment.

## Conclusions

Our study highlights the effectiveness of DNA tetrahedron (TD) structures in enhancing the delivery and, therefore, the therapeutic impact of epirubicin for treating metastatic breast cancer. The TD: Epi system demonstrated superior intracellular drug concentration and increased apoptosis rates compared to epirubicin delivered via exosomes and free epirubicin. Flow cytometry and cell viability assays indicated higher apoptosis in cells treated with the TD: Epi system, with fluorescent microscopy corroborating these results through significant live and dead cell staining. RT-qPCR analysis showed notable upregulation of pro-apoptotic genes and downregulation of the anti-apoptotic gene Bcl-2 in TD: Epi-treated cells. Results along the same lines were confirmed by Western blot analysis. While exosomes also provided improved delivery and therapeutic outcomes compared to free epirubicin, they were less effective than the TD: Epi system. These findings underscore the potential of both TD structures and exosomes in enhancing drug delivery, though TD: Epi offers a more promising approach for overcoming the challenges of poor intracellular delivery and systemic toxicity. Future studies should focus on elucidating the detailed mechanisms and evaluating the in vivo therapeutic potential of the TD: Epi system, while also exploring the complementary roles of exosomes in targeted cancer therapy.

## Ethics declaration

Not applicable

## Conflicts of interest

There are no conflicts of interest to declare.

## Disclosure statement

None declared.

## Funding

None

## Author contribution statement

S.M.A & D.B. conceptualized the manuscript. A.S., N.S., U.G., L.A.D., J.N., P.V., performed the experiments. A.S., S.S.B., wrote the main manuscript and prepared figures. S.F.A., S.M.A., reviewed the manuscript. All the authors approved the final manuscript.

## Data availability statement

Data generated in this article is already included in this manuscript.

## Acknowledgements

The authors acknowledge and extend their appreciation to the Researchers Supporting Project Number (RSPD2024R709), King Saud University, Riyadh, Saudi Arabia.

## Notes

### Competing Interest Statement

The authors have declared no competing interest.

